# Cyclin-dependent kinase 2 (Cdk2) controls phosphatase-regulated signaling and function in platelets

**DOI:** 10.1101/2020.05.31.126953

**Authors:** Paul R. Woods, Brian L. Hood, Sruti Shiva, Thomas P. Conrads, Sarah Suchko, Richard Steinman

**Affiliations:** Departments of Medicine, University of Pittsburgh School of Medicine; Departments of Hillman Cancer Center, University of Pittsburgh School of Medicine; Departments of Vascular Medicine Institute, University of Pittsburgh School of Medicine; Department of Molecular Pharmacology and Chemical Biology, University of Pittsburgh School of Medicine; The Henry M. Jackson Foundation for the Advancement of Military Medicine, Inc., Inova Women’s Service Line, Inova Health System

## Abstract

Cell cycle regulatory molecules including cyclin-dependent kinases can be recruited into non-nuclear pathways to coordinate cell cycling with the energetic state of the cell or with functions such as motility. Little is known about the role of cell cycle regulators in anucleate cells such as platelets. We report that cyclin-dependent kinase (cdk2) is robustly expressed in human platelets, is activated by thrombin and is required for platelet activation. Cdk2 activation required Src signaling downstream of the platelet thrombin receptor PAR1. Kinase-active cdk2 promoted the activation of downstream platelet kinases by phosphorylating and inactivating the catalytic subunit of protein phosphatase 1 (PP1). Erk was bound to PP1 in a complex with the PP1 regulator PPP1R12a (MYPT1) in platelets, and cdk2 inhibited the phosphatase activity of PP1 and PPP1R12a bound complexes. The requirement for cdk2 in Erk activation could be replaced by the phosphatase inhibitor calyculin if cdk2 was inhibited. Blockade of cdk2 kinase with chemical and peptide cdk2 inhibitors resulted in suppression of thrombin-induced platelet aggregation, and partially inhibited GPIIb/IIIa integrin activation as well as platelet secretion of P-Selectin and ATP. Together, these data indicate a requirement for cdk2 in platelet activation.

## Introduction

Cyclin-dependent kinases (cdks) are serine-threonine kinases best known for their role in control of the cell cycle, a function irrelevant to anucleate platelets. However, proteomic analyses have reported the presence of cdks 1-6 in platelets^1^. A cdk1 histone kinase activity has been reported in activated sheep platelets^2^ and an abstract has reported Cdk6 activity in mouse platelets^3^. While non-nuclear functions have been defined for a subset of cdk’s and/or their regulatory binding partners,^4–13^ little is known about cdk function in anucleate cells. In particular, literature on cdk activity and its regulation and potential function in human platelets has been lacking. Given that cdk’s are druggable and that several cdk-inhibitors are FDA approved or are in clinical trials, it is timely to examine the role of this class of kinases in platelets.

We hypothesized that cdk’s were recruited into platelet agonist signaling pathways. In this report, we characterize cdk2-signaling in platelets and the effects of cdk2-inhibition on platelet activity and signal transduction.

Coagulation of platelets is initiated by collagen-von Willebrand Factor (VWF) complexes and local formation of thrombin. Thrombin cleaves and activates G-protein coupled receptors PAR1 and PAR4 on human platelets. Thrombin signaling through PARs activates phospholipase C followed by calcium signaling and downstream kinase action to induce shape change, aggregation and the release from platelets of additional agonists including ADP and thromboxane A2 that promote irreversible aggregation. Feedback and feed-forward pathways amplify these signals during formation of clots. Among the kinases that are required for platelet function are Src family kinases (SFKs)^14–18^, Akt^19,20^ and Erk^16,21–24^. In cell line models, Cdk2 is activated downstream of Src and both phosphorylates and is a putative substrate of SFK Lyn^25–27^. Akt also has reported to be both a target of cdk2 and to phosphorylate it^28–30^. Similarly, cdk2 can be activated downstream of Erk^31,32^ and two reports have indicated that cdk2 and Erk are associated in protein complexes^33,34^. Whether cdk2 has functional interactions with these or other kinases in platelets is unknown.

A critical target of cdk2 in cell cycle progression is the Rb protein. Platelets have not been reported to contain Rb protein but do contain other potential cdk2 kinase targets, such as PPP1CA, the catalytic subunit of protein phosphatase 1 that undergoes inhibitory phosphorylation by cdk2 during the cell cycle^35^. Prior proteomic and SAGE analyses suggest as many as 31 candidate cdk2-interacting proteins in platelets^1^.

Our investigation characterized cdk levels in platelets and analyzed signaling upstream and downstream of cdk2 activation in platelets. To test the assumption that cdk2 activity is required for functional platelet responses to thrombin, we measured thrombin-induced aggregation and other platelet function measures in cdk2-kinase-inhibited platelets. In considering how cdk2 was integrated into platelet signaling, we focused on cdk2 effects on downstream kinases. These investigations highlighted that thrombin activation of aggregation-supportive kinases depended on inhibition of the phosphatase PP1 by cdk2 in platelets.

## Results

### Platelets express canonical cell cycle proteins

Although they are anucleate and do not undergo cell-cycle associated replication, immunoblotting showed that human platelets contain many of the regulatory proteins associated with cell cycle control including cdks 2, 4 and 6; cyclins E2, and D3, and cdk-inhibitors p21 and p27 (Figure 1). Small, potential cleavage products immunoreactive with multiple cyclin E antibodies were seen despite direct lysis of platelets in SDS loading buffer or preincubation with proteasome or protease inhibitors (not shown). We could not detect the cdk substrate protein Rb in platelets, consistent with its absence in mass spectrometry reports of the platelet proteome^1^.

**Figure 1.**
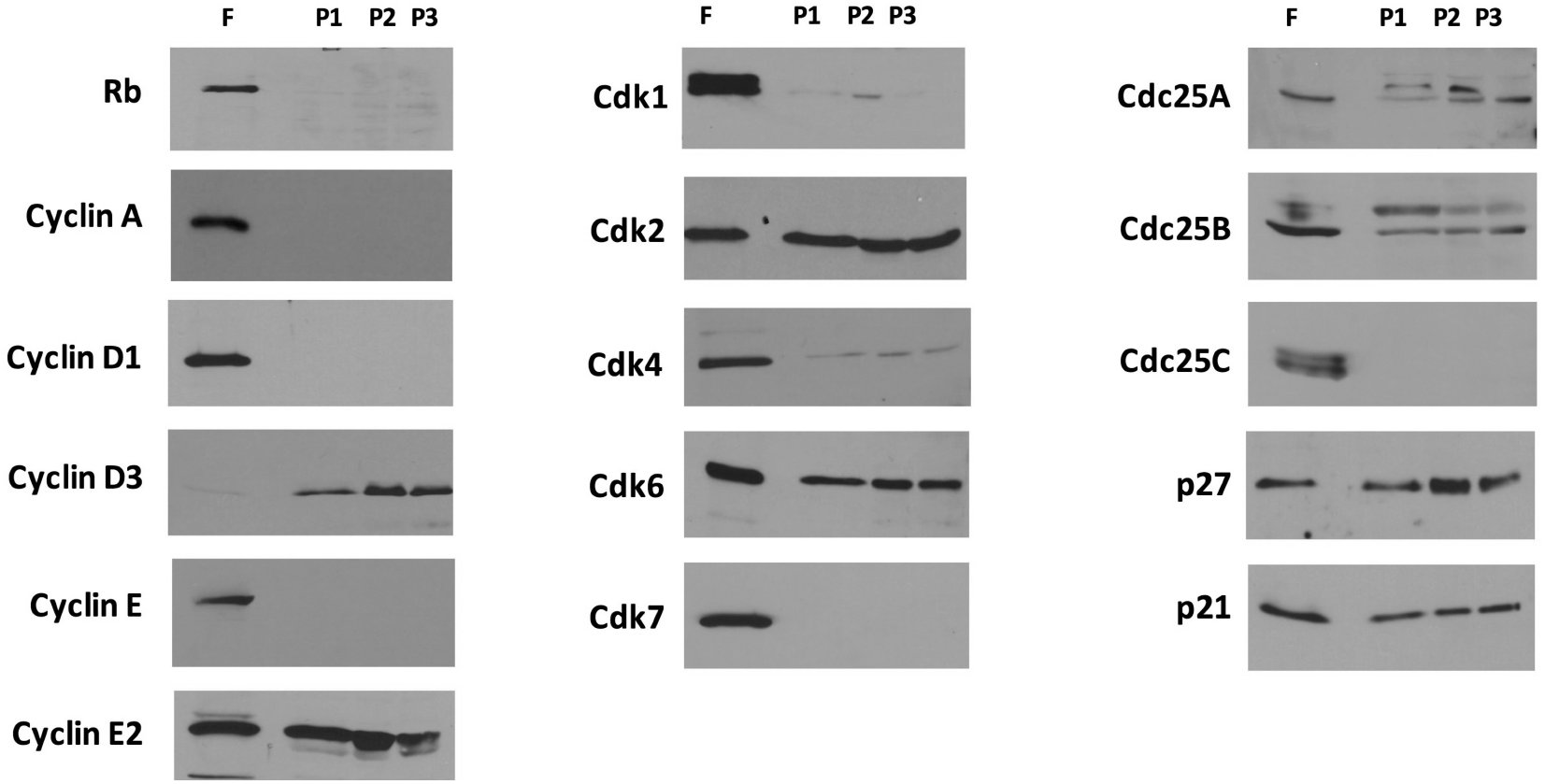
Cell cycle-related proteins in platelets. Expression of cell cycle regulatory proteins in MRC-5 human fibroblasts (F), freshly harvested human platelets prepared from peripheral blood (P1) or from freshly outdated blood bank platelets (P2, P3). Parallel immunoblots were prepared and equivalent loading verified by ponceau staining.

### Thrombin activates Cdk2 through PAR1 stimulation and Src activation

In order to determine whether cdk2 played an active role in platelet biology, rather than incidental packaging of cdk2 into platelets, we measured whether thrombin altered cdk2 activity in platelets. Figure 2A shows kinase assays demonstrating active cdk2 kinase in thrombin-stimulated but not in resting platelets. Pre-incubation of platelets with the cdk2-inhibitor roscovitine or with a cell-permeable peptide inhibitor of cdk2 (YGRKKRRQRRRGPVKRRLFG, labeled 2i)^36,37^ prevented thrombin-mediated induction of cdk2 kinase activity. In contrast, the cdk4/6 inhibitor palbociclib had no effect on thrombin activation of cdk2 (Supplemental Figure S1).

**Figure 2.**
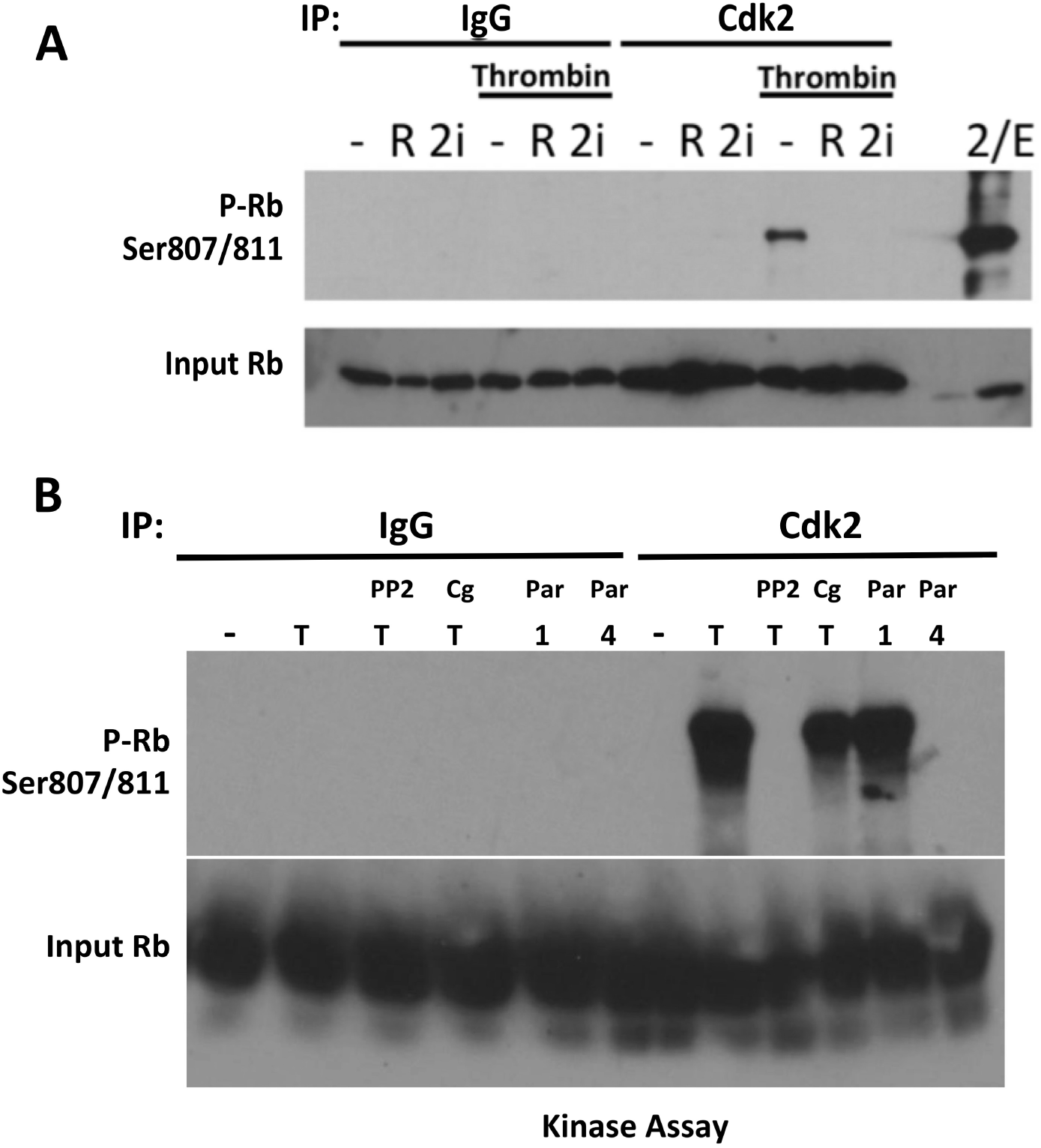
Activation of cdk2 kinase activity by thrombin. **A.** Cdk2 kinase in thrombin-exposed platelets is blocked by cdk2 inhibitors. Platelets were pre-incubated for 5 minutes with DMSO vehicle (-, 0.1 % v/v) roscovitine (R, 5 ug/ml), peptide cdk2 inhibitor (2i, 10 uM) prior to thrombin (0.5U/ml) as indicated. Positive control was recombinant cdk2/cyclinE complex (2/E). **B.** Thombin activates Cdk2 directly in a Src-dependent manner. Platelets spinning at 37° were activated for 1 minute with thrombin (T), SFLLRN (Par1, 40 uM), or AYPGKF-NH2 (Par4,200 uM) following 5’pre-incubation with PP2 (10 uM) or cangrelor (Cg, 1 uM) as indicated.

In human platelets, thrombin signals through two G-protein coupled receptors: PAR1 that is high affinity and rapid acting, and a lower-affinity PAR4 receptor. In order to determine which receptor was responsible for cdk2 activation by thrombin, kinase assays were performed on cdk2 immunoprecipitates from platelets stimulated by the PAR 1 agonist SFLLRN or the PAR4 agonist AYPGKF-NH_2_. As shown in Figure 2B, cdk2 was activated within 1 minute following PAR1 activation and not through PAR4 signaling.

To position Cdk2 in the transduction pathway of activated platelets, we determined whether blocking Src kinases signaling with the pan-Src inhibitor PP2 would prevent cdk2 activation. Thrombin activates Src kinases directly downstream of PAR1 and PAR4^15,17,38^. Src kinases contribute to platelet aggregation, secretion, integrin signaling and thromboxane A2 release (for review,^39^). Figure 2B demonstrates that PP2 prevented cdk2 kinase activation indicating that cdk2 was activated downstream of Src. Moreover, blocking cdk2 kinase with roscovitine or with peptide cdk2 inhibitor did not prevent SFK activation by thrombin (measured by autophosphorylation of tyrosine-416 in Src, Supplementary Figure S2). That finding also supports cdk2 activation downstream rather than upstream of Src in thrombin-exposed platelets.

By triggering secretion of ADP, thrombin can also activate Src kinase indirectly via ADP binding to the P2Y12 receptor^40^. In order to determine whether cdk2 activation required P2Y12 signaling, kinase activity was measured in the presence of the P2Y12 antagonist cangrelor at a concentration confirmed to block P2Y12 signaling (see also Figure 4). Thrombin activated cdk2 in the presence of cangrelor, confirming that cdk2 activation did not require secreted ADP (Figure 2B). However, blockade of P2Y12 led to partial inhibition of cdk2 activity in platelets, suggesting crosstalk between P2Y12 and PAR receptors to support cdk2 activity once initiated.

### Cdk2 is involved in the platelet response to thrombin

Figures 3A-C shows representative transmission and impedance aggregometry of platelets exposed to thrombin in the presence or absence of cdk2 inhibitors roscovitine, the cdk2-inhibitor peptide, or a type II cdk2 inhibitor, K03861(400 nM, CAS 853299-07-7)^41^. Significant suppression of aggregation by all 3 cdk2 inhibitors was evident. This was not due to compromised platelet viability; the cdk2-inhibitor roscovitine did not induce apoptosis in platelets (Supplementary Figure S3).

**Figure 3.**
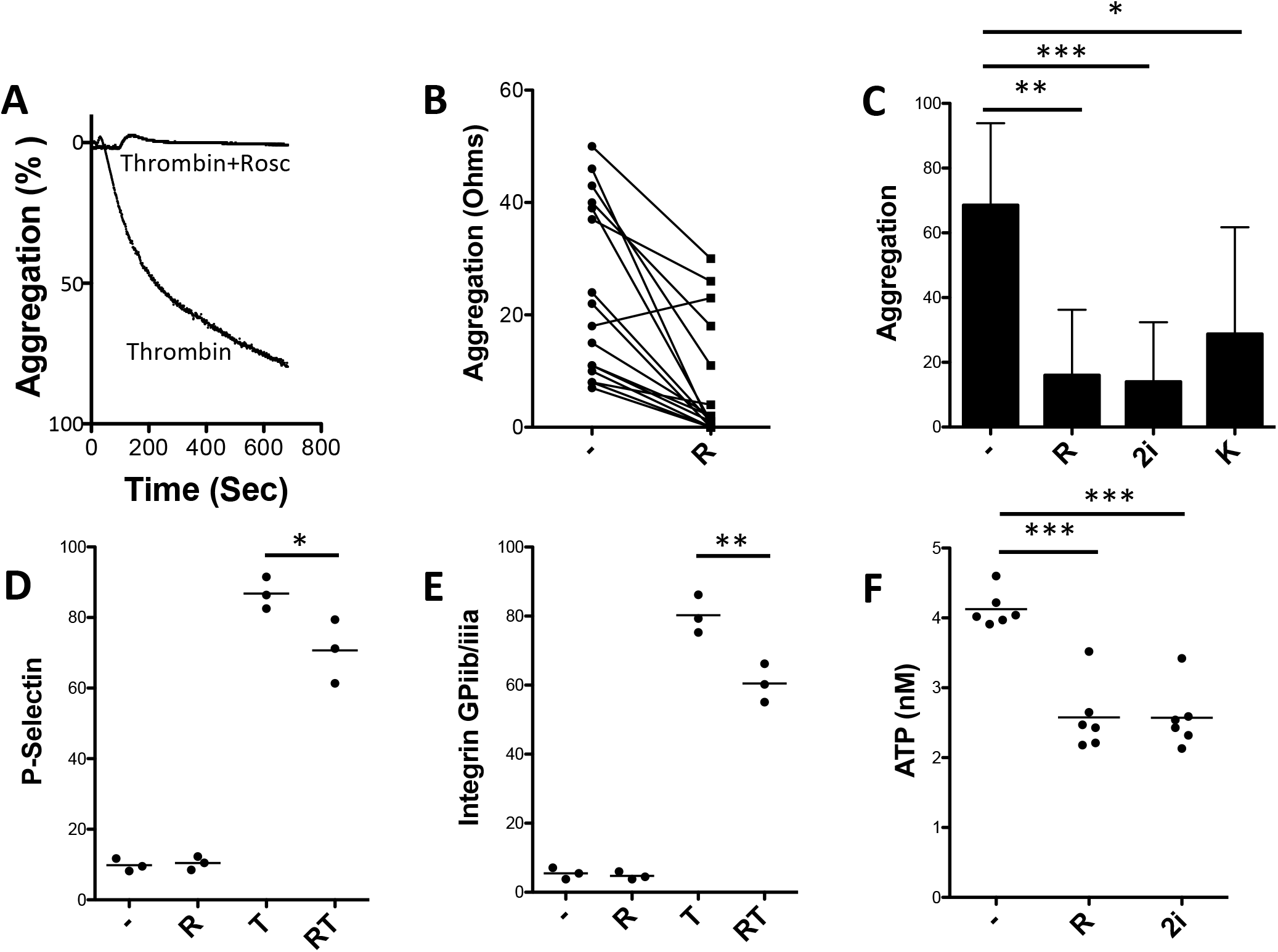
Cdk2-inhibitors suppress platelet activity. **A-C,** Cdk2 kinase requirement for aggregation. **A**. Representative transmission aggregometry of platelets exposed to vehicle (-,0.1% DMSO), Roscovitine (R, 5 ug/ml) or cdk2-inhibitory peptide (2i, 10 uM), followed by 0.25U thrombin. **B**. Impedence aggregometry of platelets exposed to vehicle or roscovitine (n=11, paired T-test, 2-sided, ****p<0.0001). **C**. Transmission aggregometry of platelets exposed to indicated cdk2-inhibitors (n=8). **D, E.**, Roscovitine suppresses surface P-Selectin expression and activation of integrin GPIIb/IIIa by thrombin. **F.** CDK2 inhibitors lower platelet secretion of ATP during 5 minute thrombin treatment. C-G, *p<.05, **p<0.01, ***p<0.001 by 1 wayANOVA, Tukey’s post-test.

In order to determine how cdk2 could be involved in platelet activation events, we measured the effect of roscovitine on thrombin-induced activation of P-Selectin, a marker secreted from alpha granules in activated platelets (Figure 3D) and the effect on activation of the platelet integrin GPIIb/IIIa (inside-out signaling, Figure 3E)^42^. Roscovitine treatment lowered expression of these activation markers, albeit not to the level of resting platelets.

Following initial platelet activation, ADP and ATP are secreted from platelet dense granules concurrent with a second wave of activation initiated by ADP binding to purinergic receptors P2Y1 and P2Y12. Figure 3F demonstrates that roscovine and the cdk2 peptide inhibitor suppressed ATP secretion, highlighting a role for cdk2 in triggering dense granule release.

### Cdk2 is required for ERK activation in platelets

Src kinases are upstream members of a platelet kinase cascade that leads to activation of AKT^15^ and the p44/42 MAPK Erk^24,43^. Given the data (Figure 2) that thrombin activated cdk2 downstream of SFK activation, we considered cdk2 as an intermediary in a pathway linking Src members to downstream kinases. To explore this possibility we measured the effect of Cdk2 inhibition on rapid (1 minute) activation of Akt and Erk by thrombin. Additional cdk2 inhibitors BMS265246^44^; and SNS-032^45^ were used to reduce the likelihood of that inhibitor effects were off target. Whereas roscovitine has activity against cdks 5, 7 and 9 in addition to cdk2^46^, BMS265246 and SNS-032 have low activity against cdk5 relative to cdk2^47,48^ and the peptide inhibitor used (2i) is specifically targeted to the cdk2 cyclin docking site.

All 5 cdk2-inhibitors blocked thrombin activation of Erk (Figure 4A, B). Roscovitine reportedly has activity against Erk so we tested the concentration used in our studies (5 ug/ml, roughly 1/2 the roscovitine IC50 v. Erk) in an independent Erk activation model in MCF7 cells^49^. Direct inhibition of Erk by roscovitine (or by the peptide cdk2 inhibitor 2i was not observed. (Figure 4C). Additionally, neither roscovitine nor peptide 2i prevented thrombin activation of pErk in endothelial cells in which, unlike in platelets, cdk2 is nuclear (Figure 4C).

**Figure 4.**
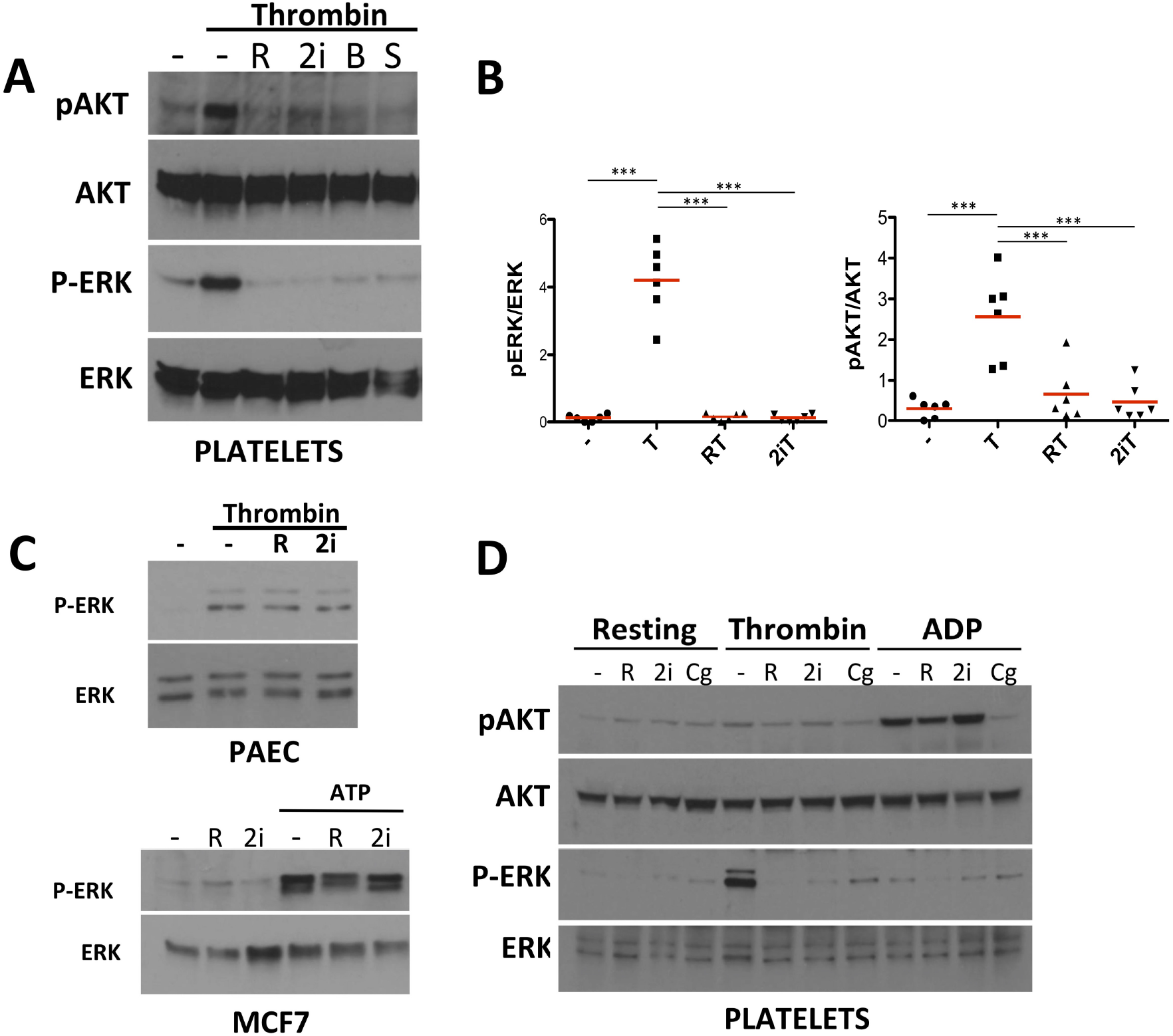
Cdk2 requirement for Erk activation in platelets. **A. B.** Inhibition of pERK in platelets preincubated 5 minutes with cdk2-inhibitors. - (vehicle, 0.1 % DMSO), R (roscovitine), 2i (peptide inhibitor), B (BMS 265246), S (SNS-032). Harvested 1 minute after thrombin (0.5U/ml); details in Methods. **C.** Cdk2-inhibitors do not block canonical Erk activation in pulmonary artery endothelial cells stimulated for 2 minutes with thrombin (0.25U/ml) or in MCF7 cells preincubated with inhibitors as in A, B, and exposed to 100 uM ATP for 2 minutes before harvest. **D.** Differential effects of Cdk2-inhibitor pre-incubation in platelets stimulated with thrombin (0.5 U/ml) or ADP (2-methylthio-ADP, 10 mM). Thrombin and ADP respectively phosphorylate Erk (Erk T202/Y204) or AKT (phospho-AKTS473) under our conditions. Lysate harvested 1 minute after agonists. Cg is cangrelor (1uM).

We considered whether cdk2 could be activating kinases directly downstream of thrombin or indirectly by promoting ADP secretion and signaling downstream of P2Y12. Thrombin activation of Erk is reportedly sensitive to the P2Y12 inhibitor cangrelor^43^.

Under our experimental conditions (1 minute exposure to agonist), Erk was phosphorylated in platelets exposed to thrombin but was not phosphorylated by exogenous ADP (Figure 4D). We confirmed that cangrelor, an inhibitor of the ADP P2Y12 receptor, suppressed thrombin activation of Erk. This result suggested that the P2Y12 signaling is necessary but not sufficient for phosphorylation of Erk. Insofar as cdk2 promotes ADP secretion, it could facilitate Erk activation by engaging P2Y12 to collaborate with thrombin signaling to Erk.

Akt was moderately activated by thrombin and more robustly by ADP. Cdk2 blockade with roscovitine or with the 2i peptide did not prevent exogenous ADP from activating pAKT, indicating that cdk2 was not required downstream of P2Y12 for AKT activation. In contrast, these Cdk2 inhibitors suppressed thrombin activation of pAKT (Figure 4B). AKT activation could require the cdk2 support of granule release and subsequent activation of P2Y12^50^ causing AKT phosphorylation.

### Activated cdk2 binds to and inhibits Protein Phosphatase 1 in thrombin-stimulated platelets

Thrombin has been reported to cause inactivation in platelets of protein phosphatase 1^51^. Cdk2 has been identified as a negative regulator of PP1 during cell cycle control, phosphorylating the PPP1CA catalytic unit at an inhibitory site *in vitro* when complexed with cyclins A and E.^35^. We explored whether cdk2 complexes in platelets had the ability to phosphorylate PPP1CA. Given that PP1 dephosphorylates and deactivates multiple kinases, this could place Cdk2 in the position of amplifying kinase cascades downstream of platelet agonists.

Our results confirmed cdk2-PPP1CA complex formation in platelets and inhibitory phosphorylation of PPP1CA at T320 site by cdk2 from activated platelets. Data in Figure 5A indicate that resting platelets expressed PPP1CA with and without inhibitory T320 phosphorylation, whereas thrombin addition fully phosphorylated the PPP1CA pool. To determine whether cdk2 bound to PPP1CA in platelets, we immunoprecipitated cdk2 in the presence and absence of thrombin. Figures 5B and 5C demonstrate that cdk2 and PPP1CA were in a complex both in activated and in resting platelets. An *in vitro* kinase assay (Figure 5D) demonstrated that immunoprecipitated cdk2 phosphorylated recombinant PPP1CA at the inhibitory Thr320 site and that this did not occur in the presence of cdk2 inhibitors or in the absence of thrombin.

**Figure 5.**
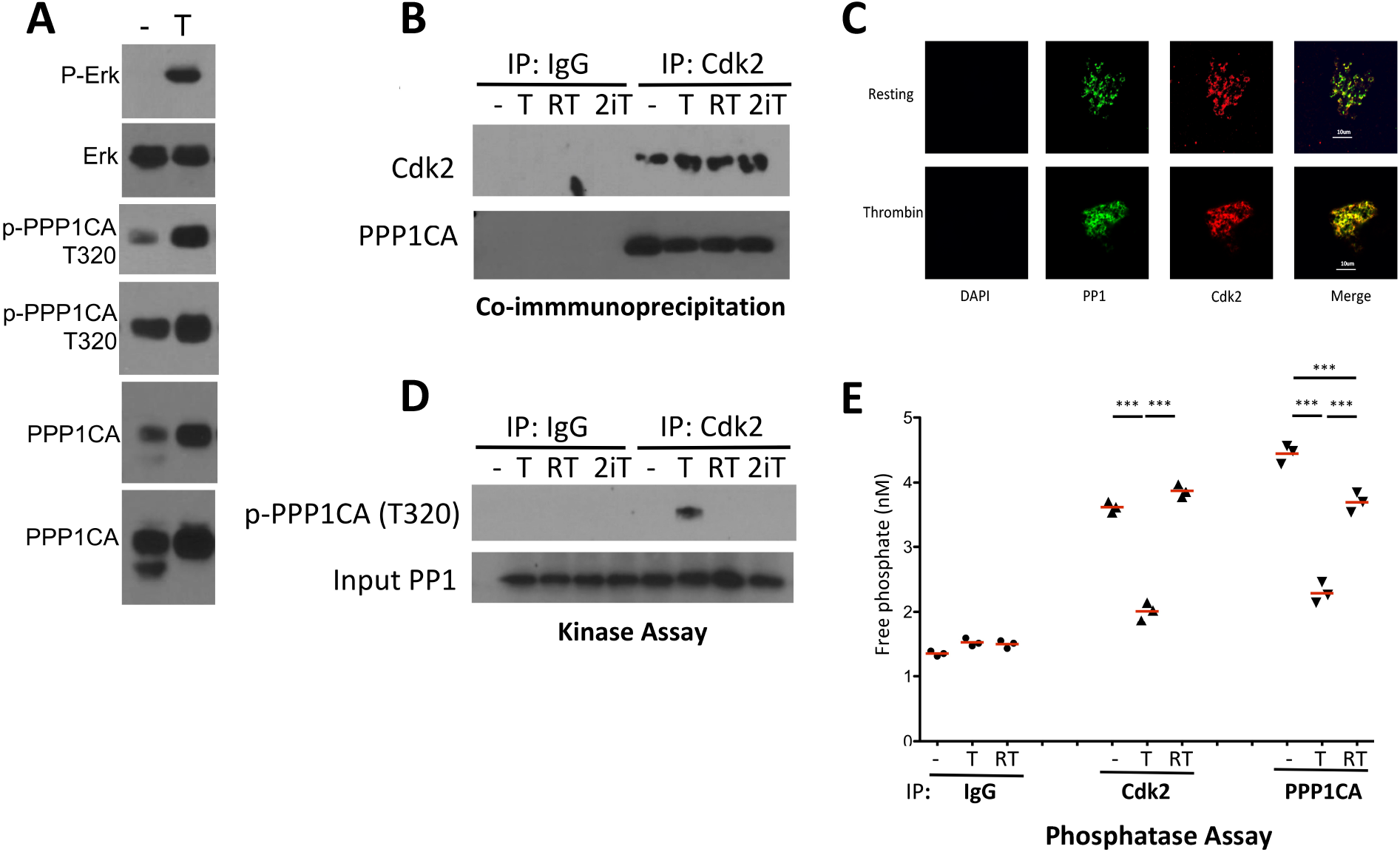
Cdk2 associates with and phosphorylates PPP1CA in platelets, inhibiting its activity. **A.** Phosphorylation of PPP1CA in thrombin activated platelets. 2 exposures of total and phospho-PPP1CA shown; the lower PPP1CA band is non-phosphorylated. **B.** Co-immunoprecipitation of PPP1CA with cdk2; - (DMSO vehicle), T (0.5 U/ml thrombin), R (roscovitine), 2i (cdk2 peptide inhibitor); 10 minute exposure to thrombin before pulldown. **C.** Immunofluorescence staining of platelets for PPP1CA (PP1) and Cdk2. Absence of DAPI signal confirmed absence of nucleated cells. 250x, bar 10 um. **D.** Kinase assay. Cdk2 (or Ig control) immunoprecipitates were incubated with recombinant PPP1CA (input, 0.5 ug/Rx) followed by WB of reaction mixture for phospho-PPP1CA. **E.** Phosphatase assay. Immunoprecipitates of platelets exposed to vehicle (-), thrombin (T) or roscovitine plus thrombin (RT) were reacted with a PP1/2a-specific phosphatase substrate and liberation of phosphate measured.

In order to confirm that cdk2 regulated PP1 activity in thrombin-activated platelets, we conducted phosphatase assays using a PP1/PP2A-specific substrate. Consistent with the binding of cdk2 to PPP1CA in both resting and activated platelets, cdk2 immunoprecipitates of resting platelets contained active PP1 phosphatase. Addition of thrombin suppressed this activity in cdk2 immunoprecipitates and also in direct immunoprecipitates of PPP1CA (Figure 5E). In order to confirm that phosphatase was specifically blocked by cdk2 activation, parallel immunoprecipitates from platelets preincubated with roscovitine prior to thrombin were analyzed. Roscovitine inhibition of cdk2 sustained phosphatase function in thrombin-exposed platelets, confirming a role for active cdk2 in inhibiting PP1. No cdk2 complexes containing PP2A could be detected and PP2A phosphatase activity was unaltered by thrombin with or without roscovitine (Supplemental Figures S4, S5). Data linked by experiment is shown in Supplemental Figure S5.

If cdk2 inhibition of PPP1CA accounts for functional outcomes of cdk2 activation and cdk2 inhibition in platelets, then chemical blockade of PPP1CA (by mimicking cdk2) should reverse the effects of cdk2 inhibitors. We used calyculin, a highly potent inhibitor of PP1 and PP2, to test this premise. As shown in Figure 6A, the addition of calyculin A (2 nM) reversed the ability of roscovitine to prevent thrombin-induced aggregation. Aggregation blockade by the peptide cdk2 inhibitor was partially reversed by calyculin A. We should note that reports that calyculin A itself prevented platelet aggregation used at least 50-fold higher calyculin concentrations^52–54^, suggesting biphasic^55^ or off-target effects; we also saw that high calyculin concentrations inhibited aggregation (Supplemental Figure S6).

**Figure 6.**
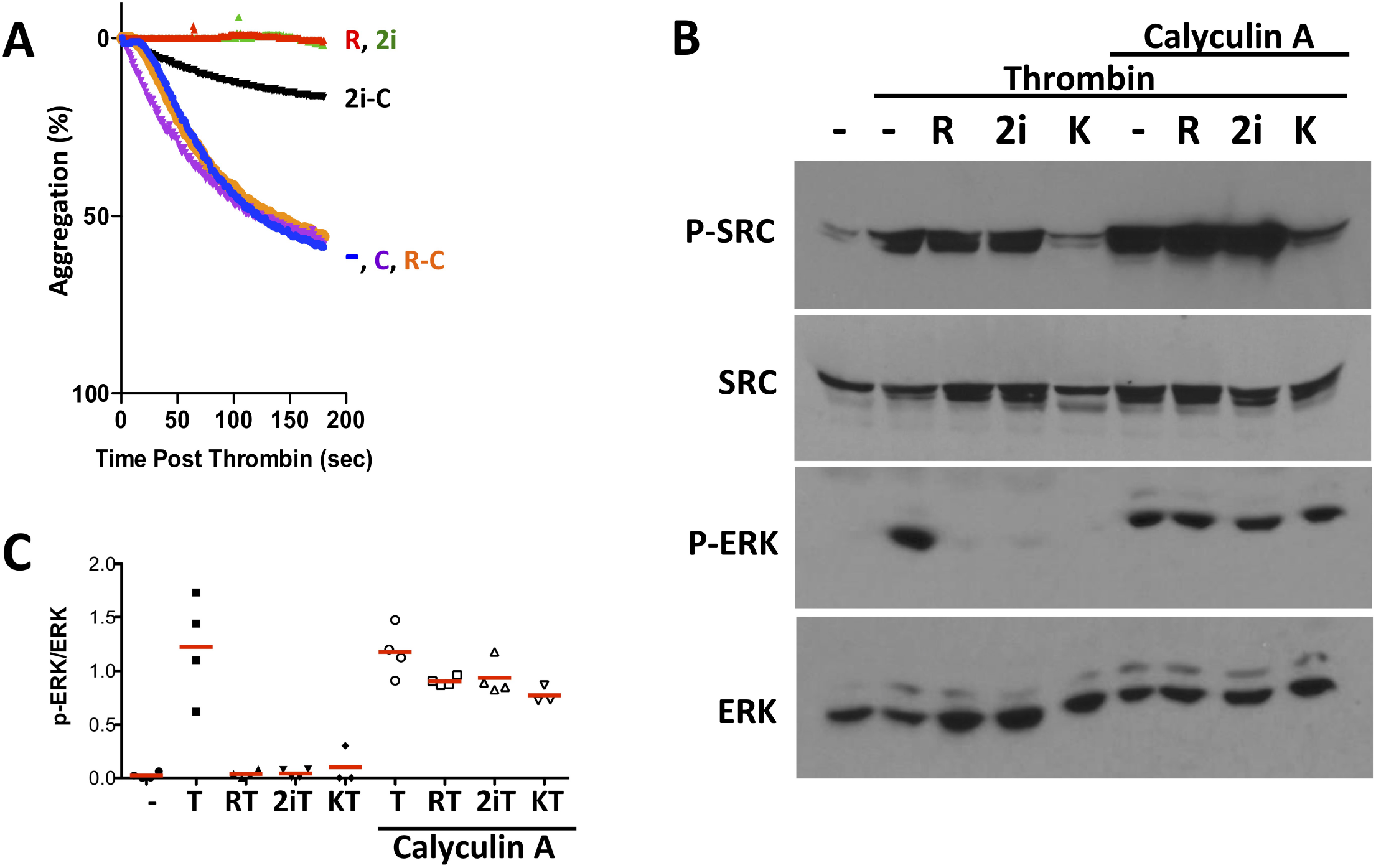
The PP1/2 inhibitor calyculin reverses effects of cdk2 inhibition in platelets. **A.** Aggregation of platelets treated with thrombin after pretreatment with vehicle (-, DMSO), roscovitine (R), peptide cdk2 inhibitor (2i) in the presence or absence of calyculin A (C, 2 nM). **B.** Immunoblot of platelets treated 1 minute with thrombin (0.5 U/ml) in the presence or absence of calyculin A and inhibitors (as in A; K is K03861 (200nM)). **C.** Quantitation of pErk in 3-4 independent experiments. Thrombin (T), inhibitors as in B. p<0.001 compared to resting platelets for thrombin alone (T) and for calyculin A-exposed platelets versus resting platelets, 1-way Anova, Dunnett’s post test.

The ability of cdk2 inhibitors to prevent Erk activation by thrombin was also reversed by calyculin A (Figures 6B, C). These results indicated that rapid stimulation of Erk phosphorylation by thrombin is not only mediated by kinase activity but also is dependent on phosphatase inactivation within the first minute of agonist activity.

### Cdk2, Erk, PP1 and PP1R12A form complexes in platelets with thrombin-responsive activity

In order to clarify whether Cdk2 contributes to phosphatase inactivation of Erk, we sought to determine whether PPP1CA was associated with Erk, and whether cdk2 co-immunoprecipitated with Erk. Newly outdated blood bank platelets were further purified (Methods) and exposed for 1 minute to thrombin while stirring at 37°. As shown in Figure 7B, Cdk2 co-immunoprecipitated Erk as well as PPP1CA.

**Figure 7.**
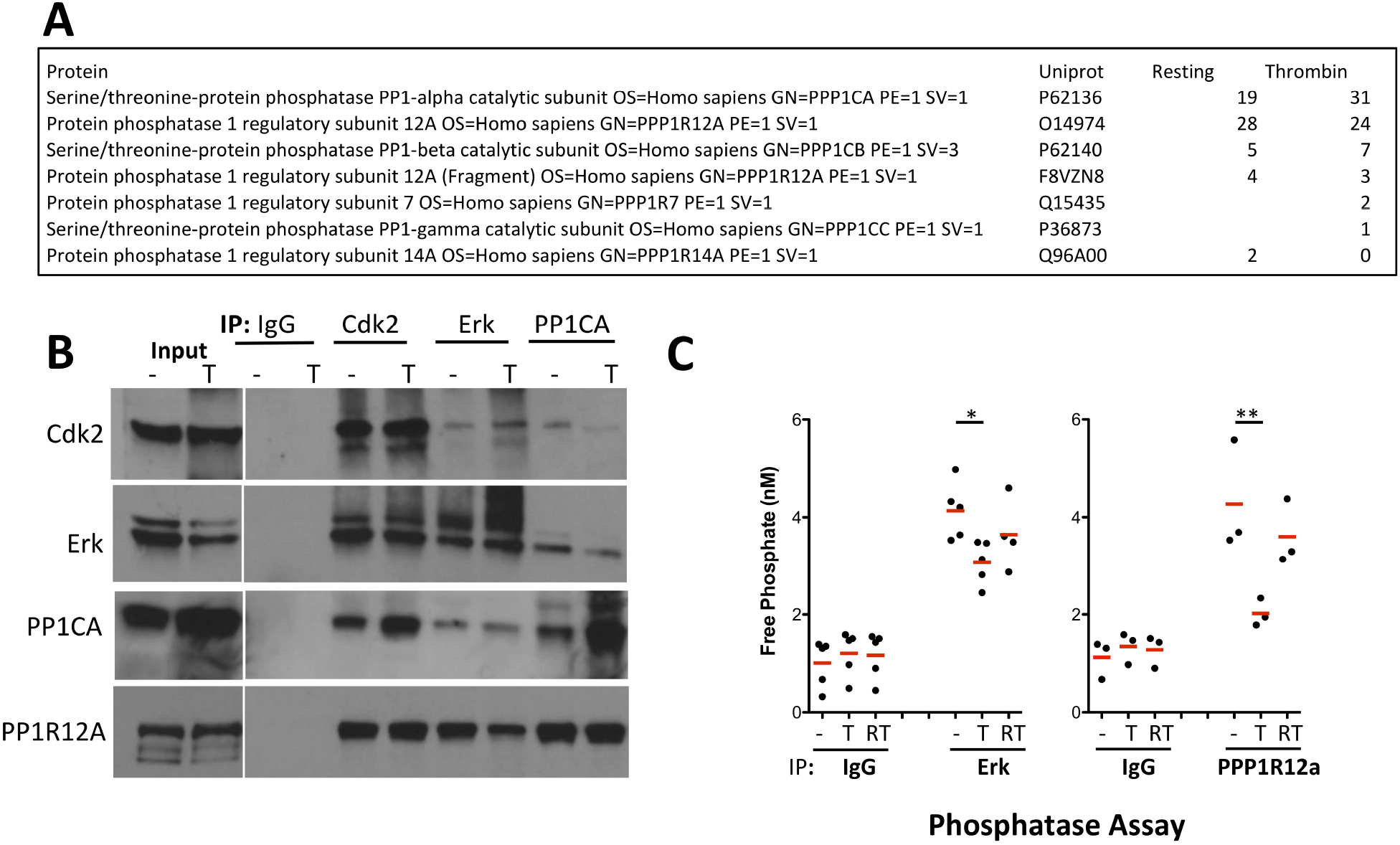
**A.** Cdk2 association with PP1 binding partners. Mass spectrometric analysis of Cdk2 immunoprecipitates (lysates pooled from 3 independent experiments) from resting or thrombin exposed platelets identified cdk2 binding to a subset of PP1 regulatory proteins. Peptide counts identified for proteins listed under each incubation condition shown; PPP1R12A was most prevalent. **B.** Co-immunoprecipitation of Cdk2, Erk, PPP1CA and PP1R12A. Lysates of platelets under resting or thrombin-exposed (T, 1 minute, 0.5 U/ml) platelets immunoprecipitated with polyclonal cdk2 or monoclonal Erk1/2 or PPP1CA antibodies as indicated were subjected to Western Blotting. Molecular weight ladder lane excised (gap); PPP1R12A input run on parallel gel. **C**. Erk and PPP1R12a-associated phosphatase activity. Immunoprecipitates of platelets exposed to vehicle (-), thrombin (T) or roscovitine plus thrombin (RT) were reacted with a PP1-specific phosphatase substrate and liberation of phosphate measured.

PP1 can be targeted to specific substrates through PP1-associated regulatory partners^56,57^. The PP1 binding partner spinophilin has been reported in neurons to target PP1 to Erk and promote its Erk dephosphorylation^58^, and other PP1 regulators and binding partners have been reported to control PP1 localization and substrate choice. In order to assess PP1 interacting partners that could deliver cdk2 to ERK, we characterized cdk2 binding partners in resting and thrombin-activated platelets by mass spectrometry analysis. Known PP1-regulators pulled down by cdk2 did not include spinophilin but the LC-MS/MS results did indicate PP1R12A as bound to cdk2 both in resting and activated platelets (Figure 7A).

PP1R12A (also known as Mypt1) binds to the PP1 catalytic unit^59^ and delivers it to myosin light chain^60,61^. PP1R12A-targeted proteins in platelets are generally unknown. As shown in Figure 7B, PPR12A was bound to Cdk2, Erk and PP1 in platelets.

We demonstrated that the PPP1R12a-directed PP1phosphatase was inhibited by cdk2 in thrombin-exposed platelets. PPP1R12a immunoprecipitates contained phosphatase activity in resting platelets, whereas thrombin suppressed this PPP1R12a-associated PP1 phosphatase (Figure 7C). Treatment of platelets with roscovitine caused higher PP1R12a-linked phosphatase activity in thrombin-treated platelets. In conjunction with results of cdk2 immunoprecipitate assays (Figure 5), this supported a role for cdk2 in inactivating PPP1R12a-bound PP1. Erk immunoprecipitates were similarly assayed for phosphatase activity. Figure 7C confirms activity of PP1 phosphatase in complex with Erk in resting platelets and suppression of Erk-associated phosphatase by thrombin. Although roscovitine addition prior to thrombin increased the phosphatase level in each experiment tested (Supplemental Figure 5), the increase of phosphatase following roscovitine treatment in thrombin-exposed platelets did not achieve significance over all experiments.

It is notable that PP1 phosphatase was in an activated form bound to Erk in resting platelets. This pre-assembly of PP1 with Erk not only positioned it for negative feedback once Erk was activated, but shows that phosphatase activity could be ongoing in the resting state to suppress subthreshold kinase signals. This is a model expressed for myosin light chain kinase in platelets^62^. In fact, an immunoprecipitation of myosin light chain confirmed associated active PP1 in resting platelets that lost its activity with thrombin treatment. As was the case for PPP1CA precipitates, loss of activity with thrombin did not occur when cdk2 was inhibited with roscovitine (Figure S5).

## Discussion

The known physiological repertoire of platelets has broadened to include not only hemostasis but also immune modulation^63,64^, augmentation of cancer growth and spread^65–67^, pathogenic sterile inflammation^68^ wound healing^69^ and bone formation. These context-dependent activities compel a new look at the intracellular signaling cascades in platelets that can drive them into functional roles linked to their microenvironment.

The functional repertoire of cell cycle proteins has also broadened from cell cycle control to include regulation of cell motility, cell survival and mitochondrial viability and activity. This study details a critical role for cdk2 in platelet function. We show that cdk2-containing complexes exist in human platelets and are integrated into functional platelet pathways. Cdk2 kinase activity was required for thrombin-mediated platelet aggregation and augmented granule release and integrin activation. This suggested that cdk2 was situated early in the signaling pathway in platelets undergoing activation by thrombin.

Interestingly, we found a significant role of cdk2 kinase in platelets to be the inactivation of protein phosphatase-1 (PP1). Reversible phosphorylation is a key process linking platelet agonists to functional outcomes. PP1 is responsible for most dephosphorylation events in cells and is highly regulated in the selection of targets and timing of phosphate removal. Our results indicated that activated cdk2 binds to and inhibits protein phosphatase 1 in thrombin-stimulated platelets. During cell cycling, Cdk2 has been identified as a negative regulator of PP1 by phosphorylating the PPP1CA catalytic unit at threonine 320, thereby preventing dephosphorylation of the Rb protein^35^. The cdk2 complexes from activated platelets similarly had the ability to phosphorylate PPP1CA at threonine 320 to block its activity.

PP1 has a complex role in the amplification and inhibition of platelet signaling. Thrombin induces dramatic relocation of this phosphatase including translocation to lipid rafts^70^ and shuttling to diverse binding partners that either sequester it or recruit it into activation pathways. These include binding of PP1 to spinophilin in activated platelets, lowering its access to myosin light chain^51^. Another report showed that PP1 dissociated from α2-integrin coincident with activating dephosphorylation of myosin light chain^71^. Thrombin induced PPP1CA shuttling from the G-protein Gβ1 to bind to and remove an inhibitory phosphate from PLCβ3, a mediator of aggregation signaling^72^. These shifts in PP1 targeting contribute to thrombin’s action: aggregation after thrombin was reduced in PPP1CA knockout mice^72^. Proper functioning of platelets clearly depends on regulated localization and activity of PP1. Multiple PP1 binding partners that target and/or regulate PP1 activity in cells^57^. Our data add cdk2 to that choreography.

Cdk2 regulation of PP1 places it at a key position in sustaining kinase cascades downstream of thrombin, given that PP1 dephosphorylates and deactivates multiple kinases. With the hundreds of PP1 substrates, it is likely that cdk2, like other PP1 partners, regulates a subset of phosphatase targets. Erk appears to be a target in a cdk2/PP1 axis because the cdk2 inhibitory peptide and roscovitine prevented Erk activation in thrombin-exposed platelets. Erk activation appeared to be blocked by cdk2 inhibitors in a PP1-dependent fashion (as indicated by calyculin rescuing activation from cdk2 blockade). Interestingly, our Western blot and phosphatase assays confirmed that functional PP1 was preassembled onto Erk prior to thrombin stimulation and that PP1 was inhibited within 1 minute of thrombin addition to the platelet.

We considered whether cdk2 was included in PP1 complexes bound to specific phosphatase targets through the actions of PP1 regulatory proteins. PP1 binds to over 200 interacting proteins^73^, and 36 PPP1CA binding proteins identified by SAGE or proteomics in platelets are listed by the Plateletweb systems biology tool^1^. Our proteomic analysis of Cdk2-binding proteins in resting or thrombin-exposed platelets identified PPP1R12a as the most prominent cdk2-bound member of the PP1 complex after PPP1CA.

PPP1R12a, cdk2 and Erk co-precipitated in both resting and thrombin-exposed platelets, suggesting a pre-formed complex that could shift its phosphatase activity depending on cdk2 activation status. Cdk2 and Erk have not previously been identified as PP1R12A partners, although one report links an inhibitory integrin polymorphism in PPP1R12a to an increase in Erk activity^74^. PPP1R12A now joins PPP1R7^75^ as an Erk-binding PP1-interacting protein.

The PP1 inhibitor calyculin rescued Erk activation in activated platelets exposed to cdk2 inhibitors, substituting for cdk2 to inhibit PP1. Our results differed from reports that used much higher calyculin concentrations in which calyculin suppressed platelet activation. It is notable that calyculin suppresses both PP1 and PP2A and to a lesser degree PP4^76^, so that these phosphatases must also be considered in interpreting calyculin results. However, the calyculin effects observed were likely attributable to PP1 (and not PP2A) inhibition because cdk2 co-immunoprecipitated of Cdk2 with PPP1CA (and not with PP2A) and because PPP1CA pulled down cdk2-sensitive PP1 phosphatase activity.

A canonical target of PP1/PPP1R12a is the myosin light chain. We have preliminary evidence that PP1 activity was associated with MLC in resting platelets and was inhibited by thrombin in a roscovitine-sensitive manner. While this could implicate cdk2: PPP1CA in the myosin light chain phosphatase, the PP1 catalytic isoform known to associate with MLC is PPP1CC (PP1-gamma catalytic subunit), not PPP1CA. Given that PPP1R12a binds cdk2 and both PP1 isoforms {Pinheiro, 2011 #7294; Hartel, 2007 #7295} and is involved in MLC dephosphorylation, cdk2 could participate in the PPP1R12a-containing complex. The mechanism underlying roscovitine modulation of MLC phosphatase remains to be clarified.

Because of the intrinsic limitations of inhibitor studies, we used multiple cdk2 inhibitors to analyze downstream effects of activated cdk2. Roscovitine can inhibit cdk2, 5, 7 and 9. Other inhibitors that mirrored roscovitine’s effect on platelets, BMS265246 and SNS-032 overlap roscovitine as cdk2, 7, and 9 inhibitors (although cdk7 appears to be absent in platelets, our data and reference^1^). We also used a cell permeable inhibitory peptide that binds the cyclin interface of cdk2 as well as a type 2 cdk2 inhibitor (K03861) that binds outside of the ATP site. Although K03861 has only been reported as a cdk2 inhibitor, our data suggest that it has partial activity against Src kinases as well.

The full range of cdk2 function in platelets may be clearer with the use of knockout models. It is of interest that cdk2 knockout mice do not have a bleeding diathesis or other signs of platelet dysfunction. This could be due to redundancy in cdk2 function with cdk4 or cdk6. Hemorrhagic edema has been reported in double cdk2 and cdk4 mice^77^; however, conditional cdk2 knockout in adult cdk4-null mice did not compromise viability^78^. Hematopoietic-specific double cdk2/4 knockouts exhibited impaired thrombopoiesis under stress^79^. Ultimately, tamoxifen-induced platelet-specific knockout of cdk2 would highlight cdk2 requirements for mouse platelets while avoiding redundancy during development. A cautionary note in reliance on such a mouse model is that PAR1 that activated cdk2 in our human studies is functionally species-specific with mice responding to thrombin through PAR3 and PAR4.

Cdk2 kinase activity was induced by thrombin through the PAR1 receptor and associated Src signaling. Src has been shown to be activated by both PAR1 and PAR4 thrombin receptors^17,80^. Positioning of Src upstream of cdk2 was supported both by the inability of thrombin to activate cdk2 kinase in the presence of the Src inhibitor PP2 and by the persistence of agonist-induced Src phosphorylation in the presence of cdk2 inhibitors.

We continue to investigate the exact mechanism of cdk2 activation. Src has been reported to activate cdk2 through inhibitory phosphorylation of p27Kip1^81^, however p27, while present in platelets, did not bind to cdk2 (Steinman, unpublished results). The Src family kinase Lyn has been reported either to inhibit or to activate cdk2^26,27^. In the canonical cell cycle, cdk2 is activated by dephosphorylation of inhibitory phospho-threonine 14 and tyrosine 15 by Cdc25, followed by activating phosphorylation at Thr160 by cyclin-activating kinase (CAK). Although we identified Cdc25A and Cdc25B in platelets, we are uncertain whether this canonical activation process was intact, because we have been unable to detect cdk2Thr160 in active cdk2 immunoprecipitates. Additionally, we could not detect the CAK kinase cdk7 in platelets. This raised the prospect that cdk2 was being activated by through a noncanonical mechanism that did not require phosphorylation of the cdk2 T-loop at T160 by CAK. Future studies will focus on noncanonical of dk2 activation in platelets.

The involvement of cdk2 in PAR1 signal transduction in platelets represents a new non-nuclear functionality for this canonical cell cycle kinase. It is notable that cdk2 did not appear to be positioned downstream of thrombin signaling in endothelial cells, where cdk2 is largely sequestered in the nucleus. The pathway uncovered here may be relevant beyond platelets to conditions (such as TGFbeta or chemotherapy)^82–84^ in which cdk2 is relocalized to the cytoplasm in PAR1-expressing, thrombin-responsive cells, enabling cdk2 to direct thrombin signaling toward dephosphorylation events.

We have showed integration of cdk2 activation into platelet signaling and function *ex vivo*. There is burgeoning interest and development of pharmacologic mediators directed at cdk2^85–87^. The priority accorded cdk2 as a pharmacologic target is timely given the possible role of cdk2 in the diverse physiology of platelets.

## Methods and Reagents

### Reagents

Cdk inhibitors were used at final concentrations as follows: R-Roscovitine (Cayman Chemical # 10009569)(CAS 186692-46-6) at 5 ug/ml; cdk2-inhibitory peptide (Apex Bio#A1107) at 10 uM^36^; K03861 (Apex Bio, #B6402)(CAS 853299-07-7) at 200 or 400 nM^88^; BMS 265246 (Cayman Chemical #19168) (CAS 582315-72-8) at 5 uM^44^; SNS-032 (Cayman Chemical #17904) (CAS 345627-80-7) at 5 uM^45^. All inhibitors were aliquoted following reconstitution at 1000x in DMSO or PBS as indicated on the manufacturer’s datasheet, for storage at −80°. Other reagents are listed in Supplementary Table 1.

### Platelet Isolation

Expired Human platelets were obtained in accordance with a protocol approved by the IRB to meet criteria for human subject exemption. The platelets suspended in platelet rich plasma were obtained from the Pittsburgh Central Blood Bank (subsequently from Vitalant, Inc.). Platelet rich plasma was centrifuged with 1uM Prostaglandin E1 at 1000 RPM for 15 minutes in a desktop Sorvall centrifuge. Supernatant were removed and placed in new falcon tube. Platelets were then pelleted at 2500 rpm for 10 minutes with no brake. The platelet pellet washed washed twice with Hepes-Tyrodes buffer (134mM NaCl, 12mM NaHCO_3_, 2.9mM KCL, 0.34mM Na_2_HPO_4_, 1mM MgCl_2_, 10mM HEPES, 5mM Glucose, 1% BSA). Platelet count was obtained via hematocytometer and adjusted to 1×10^9^/mL for all experiments. These platelets were used in lanes P1 and P2 in Figure 1 and in Figure 5, B and D.

Platelets were also isolated from whole blood obtained from healthy human donors in accordance with a protocol approved by the University of Pittsburgh Institutional Review Board. Following subject informed consent, up to 50 ml blood was collected via venipuncture without tourniquet into sodium citrate tubes (Catalog #369714, BD, Franklin Lakes, NJ). Whole blood was spun at 1000rpm for 20 minutes. Platelet rich plasma (PRP) was collected into a new tube and 1uM prostaglandin E1 was added. PRP was subsequently spun 1000 RPM for 10 minutes to remove any white blood cells. Contamination of white blood cells were determined by hematocytometer and an additional 1000 RPM spin was done if needed to remove any residual WBC. After removal of white blood cells platelets were pelleted at 2500 rpm for 10 minutes at room temperature with no brake. Platelet pellets were then gently resuspended at 1×10^9^/mL in Hepes-Tyrodes buffer with 5% BSA.

### Platelet lysate preparation and Western Blotting

Whole platelet extracts were prepared via a direct lysis of platelet pellet in 2x laemilli sample buffer (0.125 M Tris-HCL pH 6.8, 10% SDS, 20% glycerol, 5% β-mercaptoethanol, 0.03 mM Bromphenol Blue). Platelet lysates were boiled for 5 minutes in preparation for Western blot and samples were used immediately or were stored at −20 °C.

Lysates were separated using SDS-PAGE. Proteins were electrotransferred onto Nitrocellulose membranes (GE Healthcare), Ponceau stain was used to visualize equal loading. Membranes were blocked for one hour with blocking buffer (5% nonfat dry milk, 1x PBS, 0.1% Tween-20). Primary antibodies specific to unphosphorylated antibodies were diluted in blocking buffer and incubated as described in Table 2. Primary Antibodies specific to phosphorylated antibodies were diluted in 5% BSA 1x PBS, 0.1% Tween-20 as described in Table 3. Following primary antibody incubation secondary antibody was added at 1:3000 for rabbit antibodies and 1:5000 for mouse. Intensity of bands was quantitated using image J. For quantitation, band intensity per condition was normalized as percent of total signal for all assayed conditions. Ratios of phosphorylated to nonphosphorylated bands used the normalized values as input. Multiple blot exposures were done to ensure that signals were not saturated and exposures used for quantitation were in a linear range.

**Table 1.**
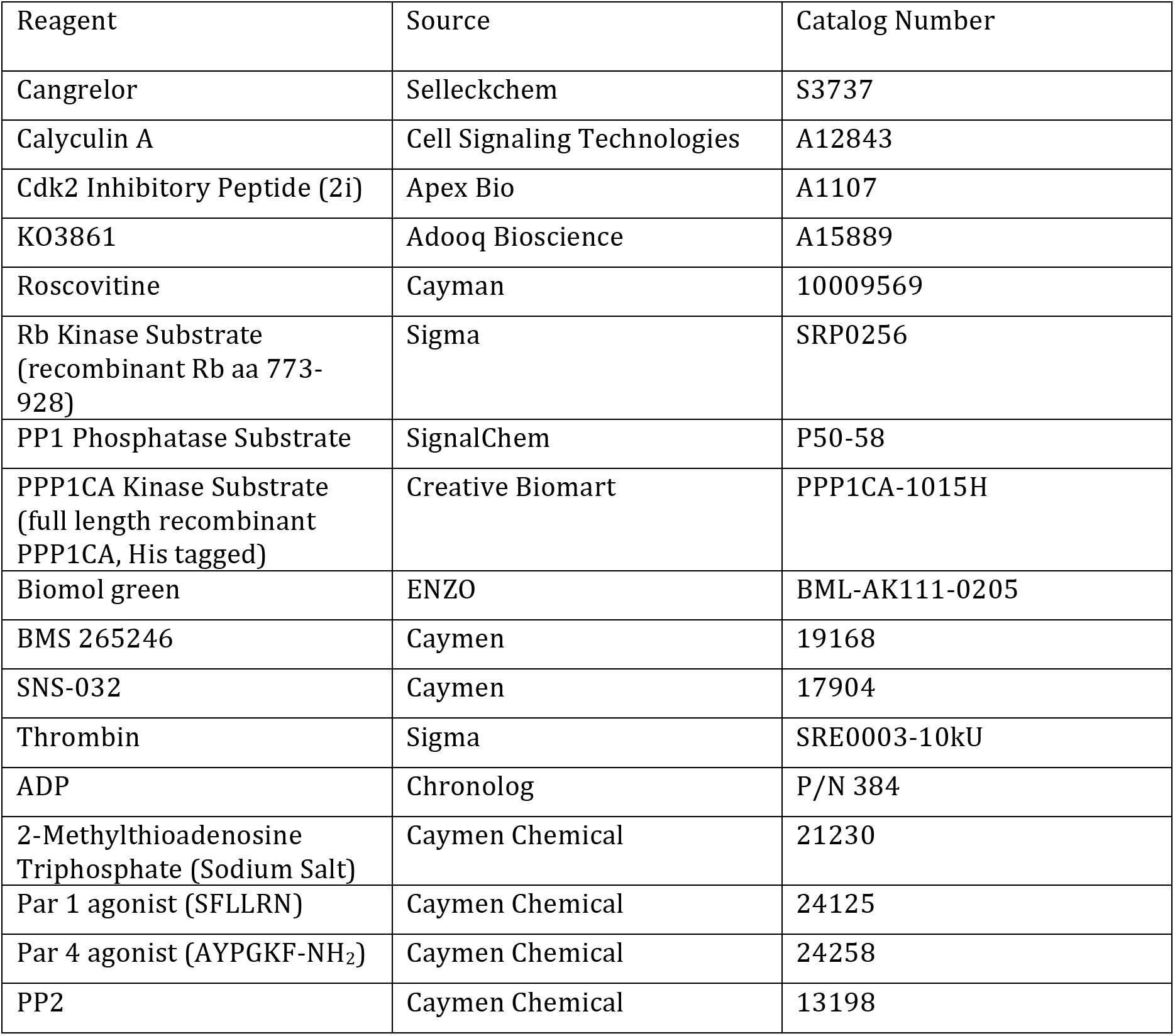
Reagents

**Table 2.**
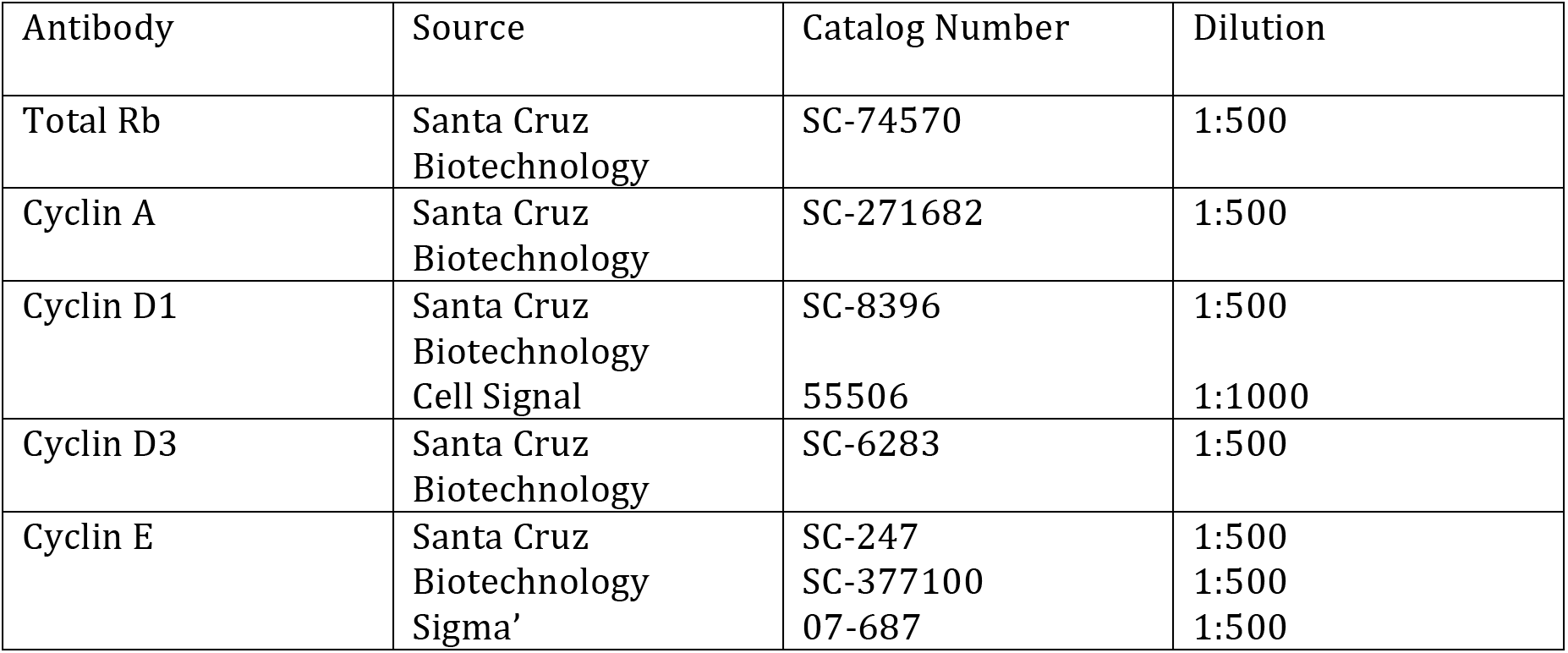
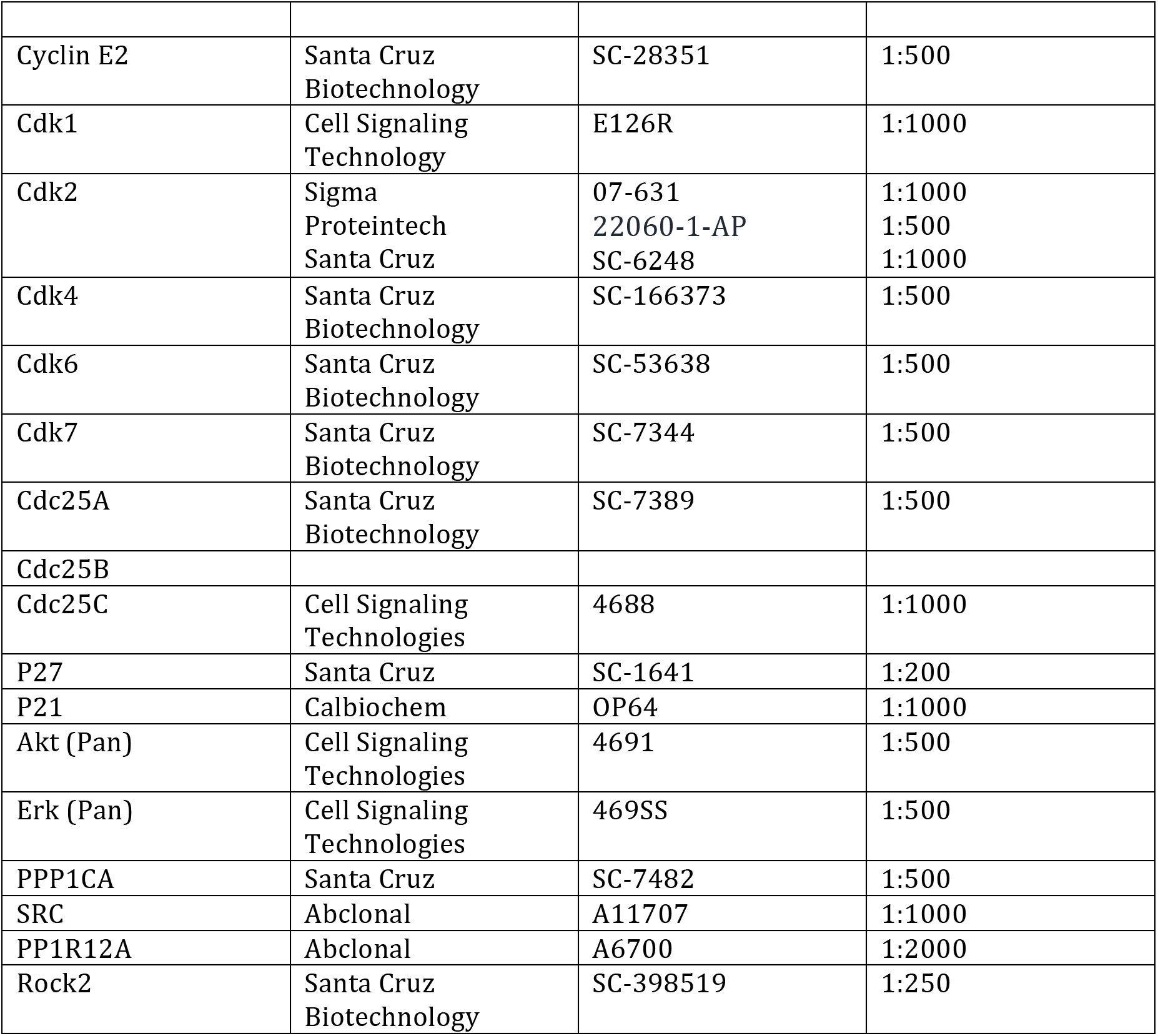
Primary Antibodies used for immunoblotting.

**Table 3.** Primary Antibodies used for immunoblotting phosphorylated proteins.

| Antibody | Source | Catalog Number | Dilution |
| --- | --- | --- | --- |
| P-Rb 807/811 | Cell Signaling<br>Technologies | 9308T | 1:500 |
| P-ERK T202/Y204 | Cell Signaling<br>Technologies | 9101S | 1:1000 |
| P-AKT S473 | Cell Signaling<br>Technologies | 4060S | 1:1000 |
| P-SRC Y416 | Cell Signaling<br>Technologies | 6943T | 1:1000 |
| P-PPP1R12A T696 | Cell Signaling<br>Technologies | 5163 | 1:1000 |
| P-PPP1R12A T853 | Cell Signaling<br>Technologies | 4563 | 1:1000 |
| P-PPP1CA T320 | Abcam | 62334 | 1:1000 |

**Table 4.** Primary antibodies used for immunoprecipation.

| Antibody | Source | Catalog Number | Amount |
| --- | --- | --- | --- |
| Rabbit IgG | Santa Cruz | SC-2040 | 5 ug |
| Rabbit Cdk2 | Sigma | 07-631 | 5 ug |
| Rabbit PPP1CA | Thermo Fisher | PA1-12376 | 5 ug |
| Mouse PPP1CA | Santa Cruz | SC-7482 | 5 ug |
| Rabbit ERK | Cell Signaling<br>Technology | 4695P | 5 ug |
| Rabbit PP1R12A | Abclonal | A6700 | 5ug |

### Phosphoprotein analysis of Human Platelets

Human platelets in Hepes-Tyrodes BSA were pelleted and resuspended in Hepes-Tyrodes without BSA at a concentration of 1×10^9^/mL. 100uL of Platelets were then spun 1200 rpm at 37°C for 5 minutes, during which they were preincubated with inhibitors or vehicle as noted in text. Following pre-incubation platelets were stimulated with either 0.25 U Thrombin (Sigma) or 10uM ADP (P/N 384, Chronolog, Havertown, PA.). Following 1 minute of stimulation 100uL of 2x laemili buffer was added to the platelets. Platelet lysates were then boiled for 5 minutes and stored −20°C for Western blot analysis of phospho-proteins.

### Immunoprecipitation

To limit signal from immunoglobulins in immunoprecipitation, primary antibodies were crosslinked to matrix using the Thermo-Fisher Pierce Co-Immunoprecipitation kit (Catalog #26149, ThermoFisher, Rockford, IL). Antibodies used are specified in Table 3. For each resin 5ug of antibody was coupled to the resin according to manufacturers instructions.

Human platelets in Hepes Tyrodes buffer with 5% BSA were pelleted by centrifugation at 2500 RPM for10 minute without brake and resuspended in Hepes Tyrodes without BSA at 1×10^9^/mL for experimental manipulations. Following 5 minute preincubation with inhibitors or other indicated reagents, agonist was added. Reactions were stopped after 1 or after 10 minutes by addition of 0.2 volumes of cold 5x lysis buffer (15mM Hepes, pH 7.4, 150mM NaCl, 5mM EDTA, 1% triton, 5ug/mL PMSF, 1mM DTT, 1mM NaF, and 1mM NaVanadate). Platelet lysates were then transferred to chilled ependorf tubes and nutated at 4° for 2 hours to complete lysis. Following nutation, debris was pelleted by spinning lysates 14,000 RPM for 10 minutes and supernatants were transferred to new chilled ependorf tubes.

To pre-clear lysates, 60uL of uncoupled resin was incubated with each platelet lysate sample of platelets for 1 hour at 4°C with nutation. Resins were discarded and the pre-cleared platelet lysates (PCL) were quantified using the Pierce BCA kit (Catalog #23225, ThermoFisher, Rockford, IL). The concentration of lysates was then equalized with lysis buffer, following which 1mg of each platelet lysate sample was incubated overnight at 4°C nutating with the indicated antibody-coupled resin. Following overnight incubation resins were washed 4 times with cold IP lysis buffer. To elute the target protein from the resin 10uL of elution buffer was added to each resin followed by centrifugation and collection. An additional 50uL of elution buffer was added to each resin, incubated for 5 minutes at room temperature and collected by centrifugation. 2X Laemmli buffer was added to each eluted sample. Samples were boiled for 5 minutes and stored at −20°C for subsequent Western blot analysis. Resins were washed with wash buffer 6x and stored at 4°C for repeated use. After a maximum of 5 uses resins were discarded and new coupled resins prepared.

### In-vitro Kinase assay

Cdk2 or IgG control were immunoprecipitated as above and reacted with recombinant retinoblastoma protein (Catalog #SRP0256, Sigma, St. Louis, MO) or recombinant protein phosphatase 1 catalytic unit alpha (PPP1CA-1015H, Creative Biomart) as kinase substrates. Immunoprecipitates were not eluted, rather Cdk2 and control IgG beads were washed twice with IP lysis buffer followed by two additional washes with chilled kinase assay buffer (50mM Hepes, pH 7.5, 10 mM MgCL2, 5mM MnCl2, 1mM DTT, 1mM NaVan, 1mM NaF). Beads then were resuspended in 50uL kinase reaction mix (kinase assay buffer +2mM ATP and 0.5ug/mL Rb or PPP1CA substrate Catalog #SRP0256, Sigma, St. Louis, MO) and incubated at 30°C for 30 minutes. The kinase reaction mix was then recovered from the beads via spin cup centrifugation. The reaction mix was mixed with laemili buffer in preparation for Western blot analysis. Kinase activity was determined by measuring phospho-Rb S807/811 or phospho-PPP1CA T320 signal compared to input recombinant substrate.

### Aggregometry and ATP measurement

Both whole blood (impedance) and light transmission aggregometry were used to analyze platelet aggregation. Platelets were stimulated with 0.25 U/mL Thrombin (Sigma) or 10 uM ADP or 2MeADP and assessed using a Chronolog Lumi-Aggregometer model 700. All experiments were performed within two hours of venipuncture. Platelets were maintained at 37° and spun at 1200 RPM during aggregometry. For ATP measurements the Chrono-Lume secretion kit was used with the aggregometer measurement program Chronolog-8 (P/N 395, Chronolog, Havertown, PA).

### Immunofluorescence

Platelet preparations at 1×10^9^/mL were spun down onto glass slides at 1000 rpm for 5 minutes using a Thermoshandon Cytospin 4 (Thermo-Fisher, Rockford, IL). Platelet cytospins were fixed in 2% paraformaldehyde, permeabilized with 0.1% Triton and blocked in PBS + 2% BSA. Platelets were incubated with primary antibodies as indicated at room temperature, washed, and incubated with alexa fluor secondary antibody (life technologies, Eugene, OR) Goat anti rabbit Alexa fluor 555 or goat anti-mouse alexa fluor 488. Following staining, platelets were washed and stained with Hoeschst (Sigma, B-2883) or DAPI to detect any cellular contamination. Slides analyzed at the University of Pittsburgh Center for Biologic Imaging using a Nikon A1 confocal microscope.

### Phosphatase Assay

Protein protein phosphatase assays were performed by immunoprecipitating indicated proteins and analyzing PP1 activity against PP1/PP2A substrate (Catalog #P50-58, Signal Chem, Richmond, BC, Canada). Immunoprecipitations were performed as described above but following wash steps 5uL of 1M Tris, pH 9.5 was added to the eluates to neutralize the pH of the elution buffer. Elutants were frozen using dry ice and stored at −80°C pending analysis.

For analysis of PP1 activity all stocks were prepared with double distilled ion free water (Catalog #10977023, Thermofisher, Rockford, IL). Eluates were thawed at 37°C and mixed to create a 1x Colormetric assay buffer (200mM Tris, pH 7.5, 5mM MgCl_2_, 1mM EGTA, 0.02% B-mercaptoethanol, 0.1mg/mL BSA) in a total volume of 75uL. 25uL of PP1/PP2A substratewas added to each sample and were incubated 30 minutes room temperature in a 96 well plate. Following incubation 100uL of Biomol green (Catalog #BML-AK111, Enzo, Lausen, Switzerlan) was added to each well. Samples were read 15 minutes following the addition of Biomol green at 262 nm using a plate reader (Model 680, Bio-rad, Hercules, CA). Sample absorbances were compared and analyzed via a standard curve with best fit line and graphed using graphpad prism 8 (San Diego, CA).

### CDK inhibition of Platelet activation via P-selectin and GpIIb/IIIa

Washed human platelets were pretreated and levels of surface P-selectin and GPIIb/IIIa measured to assess platelet activation by flow cytometery (LSR Fortessa with FASCDiva software). After treatment with agonists, platelets were labeled with APC-labeled mouse anti-human CD62P antibody (for P-selectin; BD Pharmingen; 1:1000 dilution) or FITC-labeled PAC-1 (BD Biosciences: 1:1200). Marker expression was compared between quiescent platelets and those activated with thrombin (0.25 u/L) for 30 minutes with our without roscovitine (5ug/ml) present.

### Statistics

All data are presented as mean ± standard deviation for n ≥ 3 unless stated otherwise in the figure legends. Comparisons were made using 1-way Anova with Tukey’s post-test unless otherwise stated. Analysis was conducted using GraphPad software. An alpha value of p<0.05 was considered significant.

### Mass Spectrometry Analysis

Immunoprecipitated samples were electrophoresed into a 4-15% gel (Mini-PROTEAN TGX, Bio-Rad), stained briefly with Coomassie (SimplyBlue Safe Stain, Novex, Carlsbad, CA) for protein detection. The entire gel lane was excised in a total of ten bands and processed by in-gel digestion.^89^ Individual gel band samples were resuspended in 25 mM AMB and were analyzed by LC-MS/MS on a nanoflow LC system (Easy-nLC 1000, ThermoFisher Scientific, San Jose, CA) coupled online with a Q Exactive MS (ThermoFisher Scientific) as previously described.^90^ Tandem mass spectra were searched against a UniProt human protein database (downloaded 02-02-2015; 67,865 entries) from the Universal Protein Resource (www.uniprot.org) using Mascot Daemon/Server (v.2.3.2/v.2.3, Matrix Science Inc., Boston, MA) using the automatic decoy search option. The data were searched with a precursor mass tolerance of 10 ppm and a fragment ion tolerance of 0.6 Da. Cysteine carbamidomethylation (*m/z* 57.021464) was set as a fixed modification and methionine oxidation (*m/z* 15.994915) was set as a dynamic modification. A maximum of two missed tryptic cleavages were allowed. Identified peptides (peptide spectral matches, PSMs) were filtered using an ion score cutoff of 33 resulting in a false peptide discovery rate of less than 1% for all peptides identified (determined from the decoy database search). PSMs whose sequence mapped to multiple protein isoforms were grouped as per the principle of parsimony.^91^

## Acknowledgements

This publication was made possible by grant number UL1 TR000005 (University of Pittsburgh Clinical and Translational Science Institute), the Vascular Medicine Institute, the Hemophilia Center of Western Pennsylvania, and the Institute for Transfusion Medicine.

We greatly appreciate the assistance of Christine Stehle for technical support.

## Supplementary Material

**Supplemental Figure S1.**
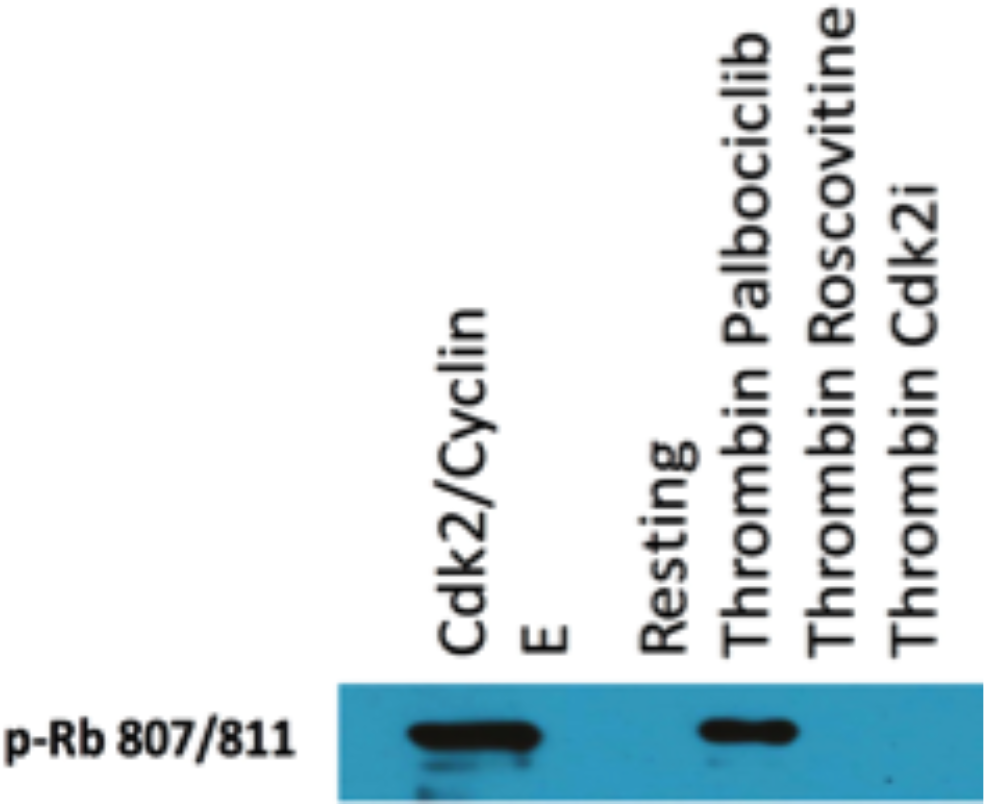
Platelets were unexposed or exposed to thrombin (0.25 U/ml) as indicated for 10 minutes following a 5 minute preincubation with DMSO vehicle (resting), palbociclib (150 ng/ml), roscovitine (5 ug/ml) or cdk2 inhibitory peptide (10 uM, cdk2i). Following immunoprecipitation with polyclonal cdk2 antibody (), a kinase assay was performed using recombinant p-RB amino acids-. A recombinant cdk2/cyclinE complex was used as a positive control, and kinase reactions were immunoblotted with anti-Phospho-Rb 807/811 antibody (Cell Signaling Technologies, #9308). Results show Rb phosphorylation by immunoprecipitated Cdk2 from thrombin exposed platelets, and blockade of phosphorylation by cdk2 (roscovitine, cdk2i) but not cdk4/6 (palbociclib) inhibitor.

**Supplemental Figure S2.**
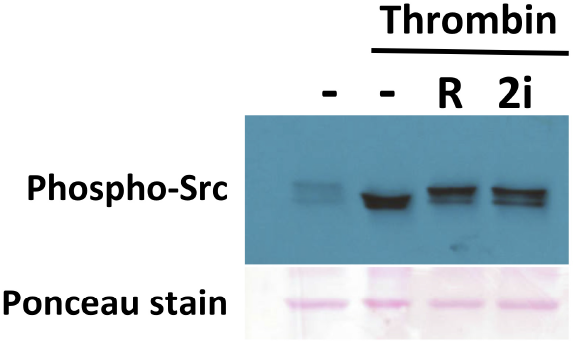
011720D. Lack of effect of cdk2 inhibitors on Src phosphorylation by thrombin in platelets. Platelets stirring at 1200 rpm at 37° were exposed for 1 minute to 0.5 U/ml thrombin and directly harvested by addition of 2X Laemmli buffer. Ponceau staining demonstrates protein loading/lane.

**Supplemental Figure S3.**
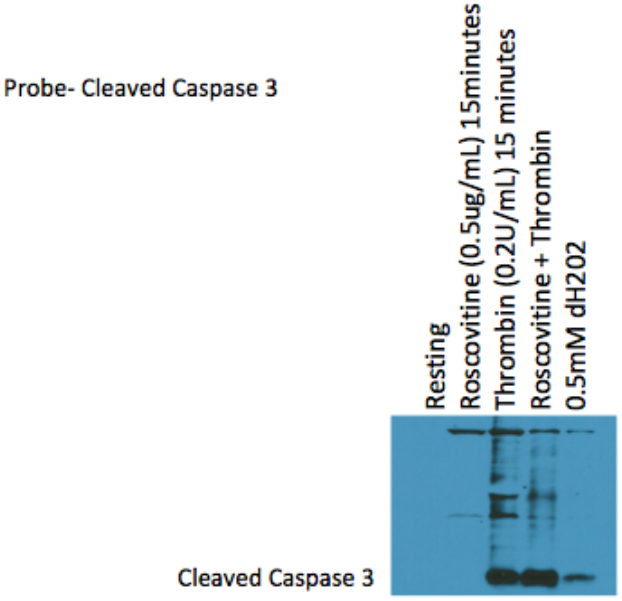
Immunoblotting platelets treated as indicated for cleaved caspase 3. Platelets were preincubated with roscovitine or with vehicle (DMSP 0.1% v/v in resting, thrombin alone) for 5 minutes following exposure to thrombin or to hydrogen peroxide (positive control) for 10 minutes. Caspase cleavage by thrombin exposure^92^ but not by roscovitine alone is evident, not does roscovitine augment apoptosis. Thrombin-mediated platelet apoptosis appears to occur through cdk2-independent pathways.

**Supplemental Figure S4.**
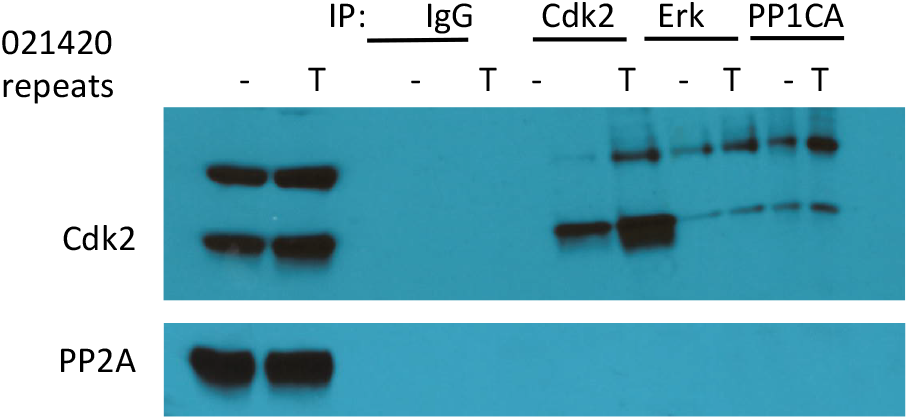
PP2A does not form a complex with cdk2 in platelets; the Sigma anti-cdk2 antibody used (07-261) detects a band at 40kD as well as cdk2 at 34kD.

**Supplemental Figure S5.**
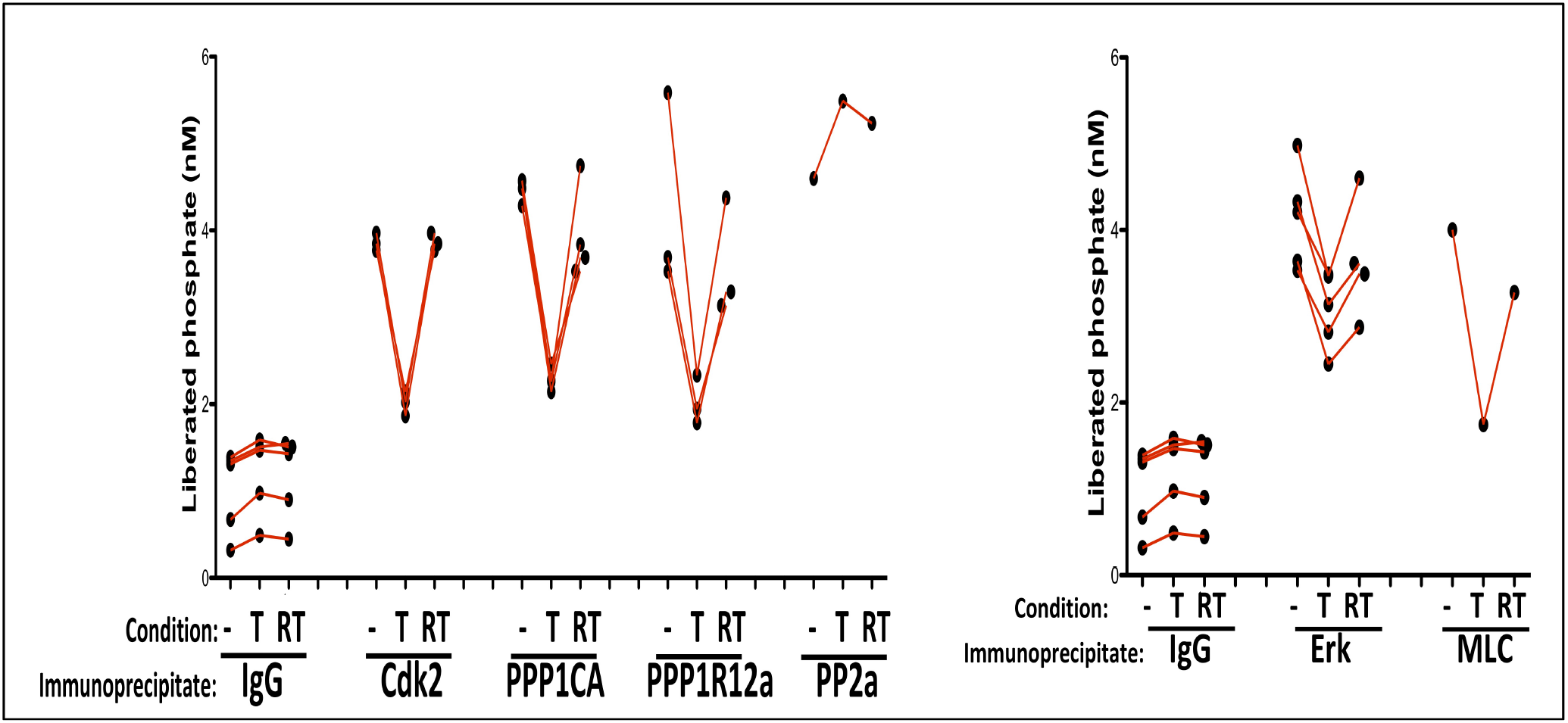
Phosphatase results shown by experiment. Cdk2 is required for thrombin to inhibit PP1 phosphatase activity. Immunoprecipitates were reacted with a PP1 phosphatase substrate and liberated phosphate measured. T, Thrombin; RT, roscovitine plus thrombin. Red line connects samples within each independent experiment. Phosphatase was active on bound proteins in resting platelets; thrombin lowers PP1 phosphatase activity and blocking cdk2 restores phosphatase activity in those complexes. In contrast, PP2a was not affected by thrombin or by the cdk2 inhibitor roscovitine.

**Supplemental Figure S6.**
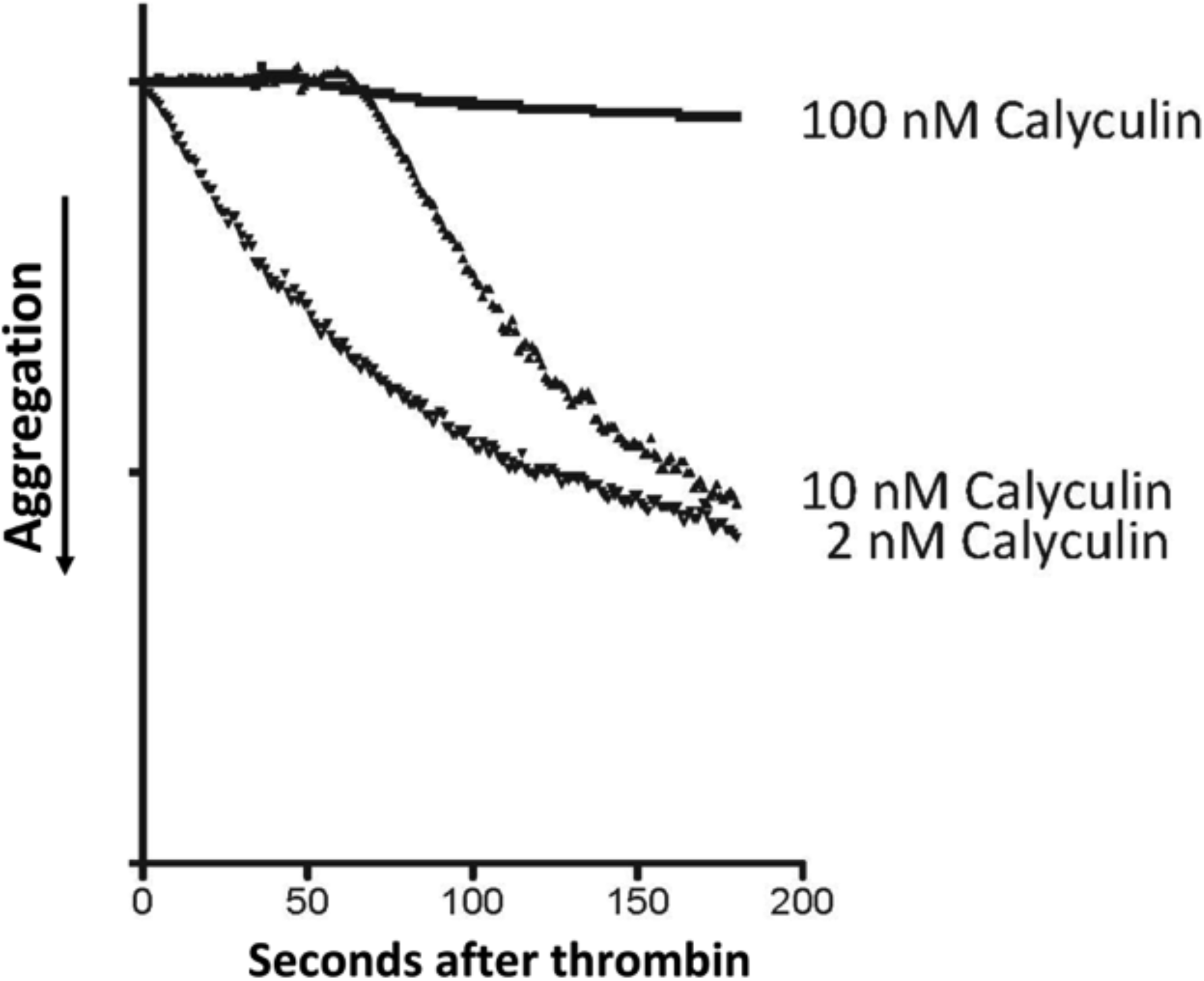
Calyculin does not inhibit platelet aggregation at the 2nM concentration used to inhibit PP1 e.g. in (Figure 6). Result confirms prior reports of aggregation inhibition at high concentrations of calyculin.

